# In vitro Kinase-to-Phosphosite database (iKiP-DB) predicts kinase activity in phosphoproteomic datasets

**DOI:** 10.1101/2022.01.13.476159

**Authors:** Tommaso Mari, Kirstin Mösbauer, Emanuel Wyler, Markus Landthaler, Christian Drosten, Matthias Selbach

## Abstract

Phosphoproteomics routinely quantifies changes in the levels of thousands of phosphorylation sites, but functional analysis of such data remains a major challenge. While databases like PhosphoSitePlus contain information about many phosphorylation sites, the vast majority of known sites are not assigned to any protein kinase. Assigning changes in the phosphoproteome to the activity of individual kinases therefore remains a key challenge.. A recent large-scale study systematically identified *in vitro* substrates for most human protein kinases. Here, we reprocessed and filtered these data to generate an *in vitro Kinase-to-Phosphosite database* (iKiP-DB). We show that iKiP-DB can accurately predict changes in kinase activity in published phosphoproteomic datasets for both well-studied and poorly characterized kinases. We apply iKiP-DB to a newly generated phosphoproteomic analysis of SARS-CoV-2 infected human lung epithelial cells and provide evidence for coronavirus-induced changes in host cell kinase activity. In summary, we show that iKiP-DB is widely applicable to facilitate the functional analysis of phosphoproteomic datasets.

## INTRODUCTION

Mass spectrometry (MS)-based proteomics provides extensive information about thousands of posttranslational modifications (PTMs)^1,2^. Phosphorylation of serine, threonine or tyrosine residues is the most extensively studied PTM in humans^3,4^, and over 200,000 phosphorylation sites have been described^5^. However, the vast majority of these sites have neither an annotated kinase nor a known biological function^6,7^. Therefore, while phosphoproteomics can now routinely quantify changes in thousands of phosphorylation sites^8,9^, the functional interpretation of these data remains a major challenge.

Computational methods to functionally annotate phosphoproteomic data can be broadly divided into two categories. The first category consists of algorithms aiming at assigning kinases to unannotated phosphosites by integrating known kinase-substrate associations with other biological features (e.g. interaction networks or co-expression profiles)^10–14^. The output of such algorithms are scores representing the likelihood of a certain site to be phosphorylated by a specific kinase. The second category of methods seeks to annotate changes in kinase activity in quantitative phosphoproteomics datasets. To this end, observed changes in phosphopeptide levels across conditions are integrated with kinase-substrate annotations^15–17^. The output of such algorithms are kinase activity profiles across conditions. Both categories of algorithms share the need for a resource representing kinase-to-phosphosite associations. The most comprehensive resource for phosphosites function is PhosphoSitePlus (PSP)^5^. However, fewer than 3% of known phosphosites have either a reported function or a known regulatory kinase in PSP^6^. Moreover, the distribution of kinase-to-phosphosite associations in PSP is skewed with a minority of well-studied kinases making up the majority of entries in the database. This bias remained essentially unchanged, despite tremendous technological developments in the field^18^.

Here, we address the challenge of predicting kinase activity in phosphoproteomic data from a different angle: instead of developing algorithms that rely on existing annotation, we expand on the knowledge of kinase-to-phosphosite annotation. To this end, we took advantage of recently published large-scale *in vitro* kinase data for over 300 human protein kinases^19^. We re-analyzed and filtered these data to compile it into an *in vitro* kinase-to-phosphosite database, or iKiP-DB. Using several published phosphoproteomics datasets of kinase activation/inhibition, we show that iKiP-DB outperforms PSP in its ability to detect changes in kinase activity. Finally, to apply iKiP-DB, we infected lung epithelial cells with SARS-CoV-2 and investigated changes in cellular kinase activity induced by the novel coronavirus.

## EXPERIMENTAL SECTION

### Analysis of *in vitro* kinase assays

*In vitro* kinase assay data from the work of Sugiyama and colleagues^19^ deposited on ProteomeXchange (dataset identifier PXD011366) was downloaded and re-analyzed using MaxQuant v1.6.10.43^20^. Each kinase assay was assigned a specific experimental group in the experimental design of MaxQuant, and, when present, replicate experiments were combined in the same experimental group. The quantification method was set to di-methyl labeling, as described in the original publication. The MS scans were searched against the Human Uniprot database (2018-07) using the Andromeda search engine. FDR was calculated based on searches on a pseudo-reverse database and set to 0.05. The search included as fixed modifications carbamidomethylation of cysteine and as variable modifications methionine oxidation, N-terminal acetylation, asparagine and glutamine deamidation as well as Phospho (STY). Trypsin/P was set as protease for in-silico digestion of the reference proteome. Phosphosites were filtered for reverse hits, contaminants and phosphosites with a localization probability lower than 0.75. Additionally, we only considered singly phosphorylated sites for building the database. Phosphopeptide intensities were log2-transformed and corrected for the intensity measured in the light di-methyl label channel, corresponding to the control experiment. As in the original publication^19^, phosphosites were considered specifically phosphorylated by a recombinant kinase when having a log-transformed ratio treatment over control higher than 1. From these kinase-substrate lists we removed all sites that were not contained in a large-scale re-analysis of 110 human phosphoproteomics experiments^7^, which was downloaded from the PRIDE repository (dataset identifier PXD012174). As a second filter, we removed all redundant phosphosites assigned to 20 or more kinases; finally, we excluded all kinase set with less than 5 annotated phosphosites. We then extracted the ±7 amino acid sequence windows around the phosphorylated site to be used as the specific identifier for each site, and combined all kinase sets into a single GMT file to be used with the ssGSEA suite to perform PTM-set enrichment analysis (PTM-SEA)^15^. Since our database is exclusively composed of kinase-substrate associations, all sites were marked as “up” sites. The database is built to work with the flanking sequence centric mode of PTM-SEA. To compare our database with PSP, we extracted the phosphosites of PTMsigDB^15^ annotated as “KINASE-PSP’’.

### Preparation of phosphoproteomics dataset for benchmarking

All datasets used for benchmarking of iKiP-DB were downloaded from the supplementary tables containing information on phosphopeptide data of the respective publications^21–25^. When not already provided, we filtered for single-phosphorylated, high confidence phosphosites (localization probability > 0.75) and calculated log2-transformed ratios of treatment(s) over control(s). We then extracted the ±7 amino acid sequence windows around the phosphorylation site (±6 for Rosales et al.) and removed duplicate entries. For each phosphoproteomics dataset, resulting sites and ratio information were exported as GCT files^15^.

### Calculation of kinase enrichments

We calculated all our kinase enrichments using the ssGSEA2.0 suite^15,26^ using the standard settings: rank sample normalization, weight for the Kolmogorov-Smirnov statistic of 0.75, area under the resulting curve as main statistic and normalized enrichment score (NES) as output score, minimum overlap between query and test set of 5 sites and 1000 permutation for p-value and NES calculations. All benchmarking datasets were analyzed using iKiP-DB and PTMsigDB^15^ (v1.9.0) separately. Since PTMsigDB contains annotations for several experimental categories, we considered exclusively sets of kinase-substrate associations (“KINASE-PSP” group). Since Rosales and colleagues only provided ±6 sequence windows for their dataset, we adapted iKiP-DB and PTMsigDB by cutting the sequences contained in the databases to ±6 and removing all duplicate entries that resulted from this cut. Analyses outputs were further processed using in-house made scripts with the R programming language. KSEA scores were calculated using in-house R scripts based on the formula described in the original publication^16^.

### SARS-CoV-2 infections of Calu-3 cells

Experiments were performed and generated in the same context as described previously by Wyler, Mösbauer, Franke and colleagues^27^. Briefly, Calu-3 cells (ATCC HTB-55) were cultivated in Dulbecco’s modified Eagle’s medium supplemented with 10% heat-inactivated fetal calf serum, 1% non-essential amino acids, 1% L-glutamine and 1% sodium pyruvate in a 5% CO2 atmosphere at 37 °C. SARS-CoV-2 (Patient isolate, BetaCoV/Munich/BavPat1/2020|EPI_ISL_406862) was grown on Vero E6 cells and concentrated using Vivaspin® 20 concentrators (Sartorius Stedim Biotech). Virus stocks were stored at -80°C, diluted in OptiPro serum-free medium supplemented with 0.5% gelatine and phosphate-buffered saline. Mock infected controls were generated with cells inoculated with cell culture supernatants from uninfected Vero cells in accordance with virus stock preparation. For the infection experiments, Calu-3 cells were seeded at 6 × 10^5^ cells/mL cells/mL and after 24 hours cells were infected with SARS-CoV-2 at an MOI of 0.33, or with Vero E6 medium as control. Samples were harvested in three biological replicates after 4, 8 and 12 hours with pre-warmed trypsin for 3 mins at 37 °C, after removal of the cell culture media. Mock infected controls were similarly harvested in biological triplicates after 4 and 12 hours. Samples were lysed and inactivated through boiling in SDS sample buffer. All infection experiments were carried out under biosafety level three conditions with enhanced respiratory personal protection equipment.

### (Phospho)-proteomics sample preparation

Proteomics samples were prepared combining SP3 sample preparation^28^ and TMT labelling^29^ using TMTpro reagents^30^. Briefly, lysates were pre-cleared via centrifugation, then reduced and alkylated with DTT and iodoacetamide, respectively. Proteins were then incubated with a 1:1 mix of SeraMag beads A and B at a 10:1 weight:weight bead:protein ratio. Protein binding to the hydrophilic beads was induced by adding ACN and washed with 80% EtOH to remove contaminants. Protein digestion was performed on beads in 50mM HEPES pH8, with Trypsin and LysC at a 1:50 protein:enzyme ratio for approximately 16 hours. After digestion, peptides were quantified via BCA assay and directly labelled with TMTpro reagents (Thermo Fisher Scientific; product number A44520, lot number UL297970) following the manufacturer’s protocol. Samples were randomly assigned to a TMT channel as follows: CoV2 4hrs repA -> 128C, CoV2 4hrs repB -> 129N, CoV2 4hrs repC -> 133N, CoV2 8hrs repA -> 133C, CoV2 8hrs repB -> 127N, CoV2 8hrs repC -> 131C, CoV2 12hrs repA -> 130C, CoV2 12hrs repB -> 130N, CoV2 12hrs repC -> 131N, mock 4hrs repA -> 132N, mock 4hrs repB -> 128N, mock 4hrs repC -> 132C, mock 12hrs repA -> 129C, mock 12hrs repB -> 126, mock 12hrs repC -> 127C. The last TMTpro channel (134) was comprised of a supermix of the other samples. Labelled peptides were pooled in equal amounts and desalted with SepPak columns (Waters) and the resulting peptide mixture was offline separated with high pH reverse phase fractionation^29,31^ on an Dionex 3000 system (Thermo Fisher Scientific) and a XBridge Peptide BEH C18 (130Å, 3.5 µm; 2.1 mm x 250 mm) column (Waters). Peptides were resuspended in high pH buffer A (5mM ammonium formate, 2% ACN) and separated on a multi-step gradient from 0 to 60% high pH buffer B (5mM ammonium formate, 90% ACN) 96 minutes long and collected in 96 fractions (1 fraction/min). The fractions were automatically pooled during collections, where each xth fraction was combined with the x+25th, x+49th, x+73th fraction for a total of 24 fractions. Of each pooled fraction approximately 1µg of peptide was subjected to mass spectrometric (MS) analysis for total proteome measurement. The remaining peptides were further pooled into 12 fractions and used as input for a phosphopeptide enrichment via immobilized metal affinity chromatography (IMAC), which was performed by the Bravo Automated Liquid Handling Platform (Agilent) with AssayMAP Fe(III)-NTA cartridges.

### LC-MS/MS Analysis

Proteome and phosphoproteome fractions were online-fractionated on a EASY-nLC 1200 and acquired on an Exploris 480 mass spectrometer (Thermo Fisher Scientific) operated on profile-centroid mode, as previously described^32^. Peptide separation was achieved on a fused silica, 25 cm long column packed in-house with C18-AQ 1.9 µm beads kept at a temperature of 45°C. Mobile phase A consisted of 0.1% FA and 3% ACN in water, while mobile phase B consisted of 0.1% FA and 90% ACN. After column equilibration peptides resuspended in buffer A were separated with a 250 µl/min flow on a 110 minutes gradient:mobile phase B increased from 4% to 30% in the first 88 minutes, followed by an increase to 60% in the following 10 minutes, to then reach 90% in one minute, which was held for 5 minutes. The MS was operated in data dependent acquisition, with MS1 scans from 350 to 1500 m/z acquired at a resolution of 60,000, maximum injection time (IT) of 10 ms and an automatic gain control (AGC) target value of 3e6. The 20 most intense precursor ion peaks with charges from +2 to +6 were selected for fragmentation, unless present in the dynamic exclusion list (30 s). Precursor ions were selected with an isolation window of 0.7 m/z, fragmented in an HCD cell with a normalized collision energy of 30% and analyzed in the detector with a resolution of 45,000 m/z, AGC target value of 1e5, maximum injection time of 86 ms or 240 ms for total proteome and phosphoproteome analysis respectively.

### (Phospho)-proteomics data analysis

RAW files were analyzed using MaxQuant^20^ v1.6.10.43, where TMTpro was manually included as a fixed modification and quantification method. Correction factors for each TMT channel as provided by the vendor were added to account for channel spillage and minimum reporter precursor intensity fraction was set to 0.5. The MS scans were searched against human and SARS-CoV-2 Uniprot databases (Jan 2020 and Apr 2020 respectively. SARS-CoV-2 database was modified to include the D614G mutation on the Spike protein) using the Andromeda search engine. FDR was calculated based on searches on a pseudo-reverse database and set to 0.05. The search included as fixed modifications carbamidomethylation of cysteine and as variable modifications methionine oxidation, N-terminal acetylation, and asparagine and glutamine deamidation. Trypsin/P was set as protease for in-silico digestion of the proteome database. Total proteome and IMAC-enriched phosphopeptides samples were analyzed in the same MaxQuant run in separate parameter groups with the same settings, except for the IMAC-enriched samples also Phospho (STY) was added as variable modification. Protein contaminants, hits in the reverse database, only identified by modified site and identified by less than two peptides of which less than one unique were removed from the ProteinGroups result table. Phosphosites were filtered by hits in the reverse database and potential contaminants. Additionally, only sites with localization probability higher than 50% were considered for further analysis. TMT reporter ion intensities for each sample were then log2-transformed and median normalized. Ratio of treatment over control were calculated with the matching mock infection for the 4 and 12 hours time point, while the 8 hours time point was corrected with the 4 hours control. Significantly regulated proteins at 12 hours post infection were calculated with a Student t-test with Benjamini-Hochberg corrected p-values. Significant cut-off was set at 10% FDR, while we used a data driven approach for a fold-change cut-off. Specifically, for each side of the distribution of fold-changes (higher or lower than zero), the function describing the density of points on the x-axis was calculated. The cut-off was set based on the x value for which 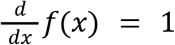 for *x* ≥ 0 and 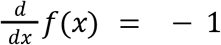 for *x* < 0. Phosphoproteomics data for PTM-SEA analysis was prepared as described above for the benchmarking datasets. All statistical analysis was done with the R programming language (v3.6.6).

### Data availability

The mass spectrometry proteomics data have been deposited to the ProteomeXchange Consortium via the PRIDE^33^ partner repository with the dataset identifier PXD030395. Prior to publication, the data can be accessed with the following credentials: **Username:**; **Password:** nLLVTHZZ

## RESULTS AND DISCUSSION

### Deriving iKiP-DB from *in vitro* phosphorylation assays

To develop our database, we took advantage of a recent large-scale discovery study of substrates of the human kinome^19^. In this study, de-phosphorylated HeLa cell lysate was incubated with 385 recombinantly expressed human kinases to perform *in vitro* kinase assays. After proteolytic digestion and phosphopeptide enrichment, the sites phosphorylated by each kinase were identified via MS (Figure 1A). Intrigued by this study, we retrieved the raw mass spectrometry data (ProteomeXchange dataset identifier PXD011366) and re-analyzed it with MaxQuant^20^. In total, we obtained 159,618 kinase-to-phosphosites associations for 17,225 unique phosphosites, localized on 4,032 distinct protein groups (Figure 1A). *In vitro* kinase assays can induce phosphorylation of sites that are not phosphorylated under physiological conditions. Therefore, we first filtered the data using a catalogue of 112 manually curated datasets of phospho-enriched proteins from 104 different human cell types or tissues^7^. More than half of the *in vitro* sites were not observed in any cell or tissue dataset and therefore excluded (Figure 1B). As a second filter, we excluded all sites assigned to 20 or more kinases, since we reasoned that these highly redundant sites are not well suited to distinguish between different kinases (Figure 1C). Finally, we removed all kinases with less than five phosphosites since such a small number would not provide sufficient information to compute robust enrichment scores.

**Figure 1.**
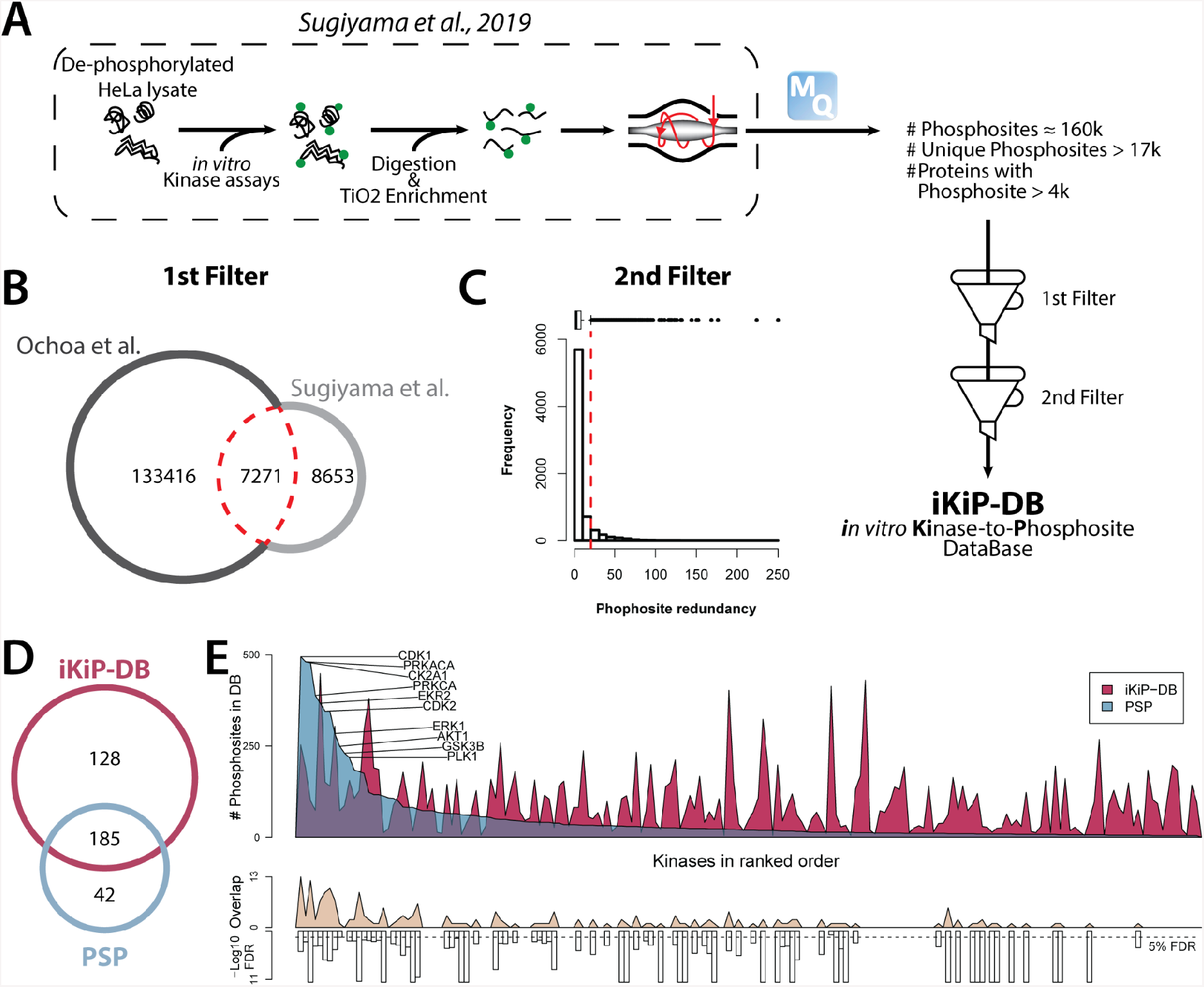
Generating iKiP-DB. (A) Schematic representation of how the kinase assay data were produced, together with the output of our re-analysis and the major steps to derive iKiP-DB. (B) Venn diagram representing the overlap between the sites identified by Sugiyama et al. and the collection of human phosphosites from the work of Ochoa and colleagues. The red line indicates which sites were selected by this first filtering step. (C) Frequency distribution of phosphosites redundancy, as in the number of kinases each phosphosite was assigned to. The red line indicates the cut-off for the second filter we applied to obtain iKiP-DB. (D) Venn diagram showing the overlap of kinases annotated in iKiP-DB and PhosphositePlus (PSP). (E) Kinases present in PSP and iKiP-DB ranked based on the number of sites annotated in PSP. Above, we indicated the number of sites in each database and the names of the kinases with most sites in PSP (higher than 200). Below, is depicted the size of the overlap between the two databases and the significance of the overlap (when present) based on an hypergeometric test with BH adjusted p-values. The dashed line indicates the 5% FDR cut-off.

After filtering, we obtained an *in vitro* kinase-to-phosphosite database (or iKiP-DB) for 313 kinases (Supplemental Table S1). iKiP-DB provides kinase-substrate associations for 128 kinases not previously annotated in PSP (Figure 1D). When looking at the 185 kinases shared between both databases, PSP showed a strong bias towards few well-characterized kinases (e.g. CDK1, PRKCA, CK2A1) while most other kinases have only a small number of assigned sites. In contrast, kinase-substrate associations in iKiP-DB do not display such a bias and greatly extend the annotations for many kinases (Figure 1E, upper panel). Nevertheless, although PSP and IKiP-DB have been derived in a completely independent fashion, we observed a significant overlap between the number of phosphosites assigned to the same kinases in both databases (Figure 1E, lower panel).

### iKiP-DB predicts the activation of EGFR and downstream pathway

To assess if iKiP-DB can predict kinase activity in phosphoproteomic data, we first turned to epidermal growth factor receptor (EGFR) signalling^34^. To this end, we took advantage of a recent phosphoproteomic dataset^21^. In this study, the authors treated HeLa cells with EGF or TGFɑ over a time course, enriched for phosphopeptides via combined phospho-tyrosine and titanium-dioxide enrichment, and measured the resulting phosphoproteome by MS (Figure 2A). This dataset provides an excellent benchmark for our database since i) EGFR/PI3K/AKT pathway is well studied and characterized^34^; ii) the time course provides a longitudinal dimension to evaluate kinases activity annotations over time; iii) EGF and TGFɑ act on the EGFR receptor producing similar but not identical outcomes^21,34^. To calculate kinase enrichments, we employed the PTM-SEA algorithm^15^, a method recently developed to calculate enrichment of specific PTM sets in MS data. Similarly to a gene set enrichment analysis (GSEA)^26^, PTM-SEA computes a rank-based statistic for sites present in an annotated PTM set within the overall distribution of all ranked sites (ordered by their abundance ratio or intensity). For comparison, we computed kinase enrichments with the same algorithm but using a database of kinase sites in the PSP collection of PTMsigDB^15^ (Figure 2B, S1A and Supplemental Table S2). The two databases produced very similar predictions for EGFR and the two downstream kinases AKT1 and RPS6KA1 (also known as p90RSK or MAPKAPK1)^21,34^. In line with previous findings, our analysis reveals that EGFR is rapidly activated upon stimulation with both EGF and TGFɑ, followed by a decrease in activity caused by receptor internalization^21^. Both analyses indicate a stronger re-activation of EGFR upon TGFɑ treatment, which has been linked to the more efficient recycling of the receptor to the plasma membrane^21^. Of note, both PSP and iKiP-DB predicted EGFR at time point 1 min as the most activated kinase throughout the treatment, confirming the specificity of the predictions (Supplemental Table S2). The relevant phosphosites for EGFR from PSP and iKiP-DB showed similar trends across the dataset (Figure 2C). Additionally, the delayed and more sustained activation of AKT1 and RPS6KA1 is in line with the more downstream position of these kinases in the EGFR/PI3K/AKT pathway^35,36^ (Figure 2B). To ensure robustness of iKiP-DB across different algorithms to calculate kinase enrichments, we also used kinase-substrate enrichment analysis (KSEA)^16^ and calculated KSEA scores and statistics, obtaining overall similar results (Figure S1B, Supplemental Table S2). In summary, these data show that iKiP-DB can be used to predict the activation status of a well-studied kinase as well as a manually curated database.

**Figure 2.**
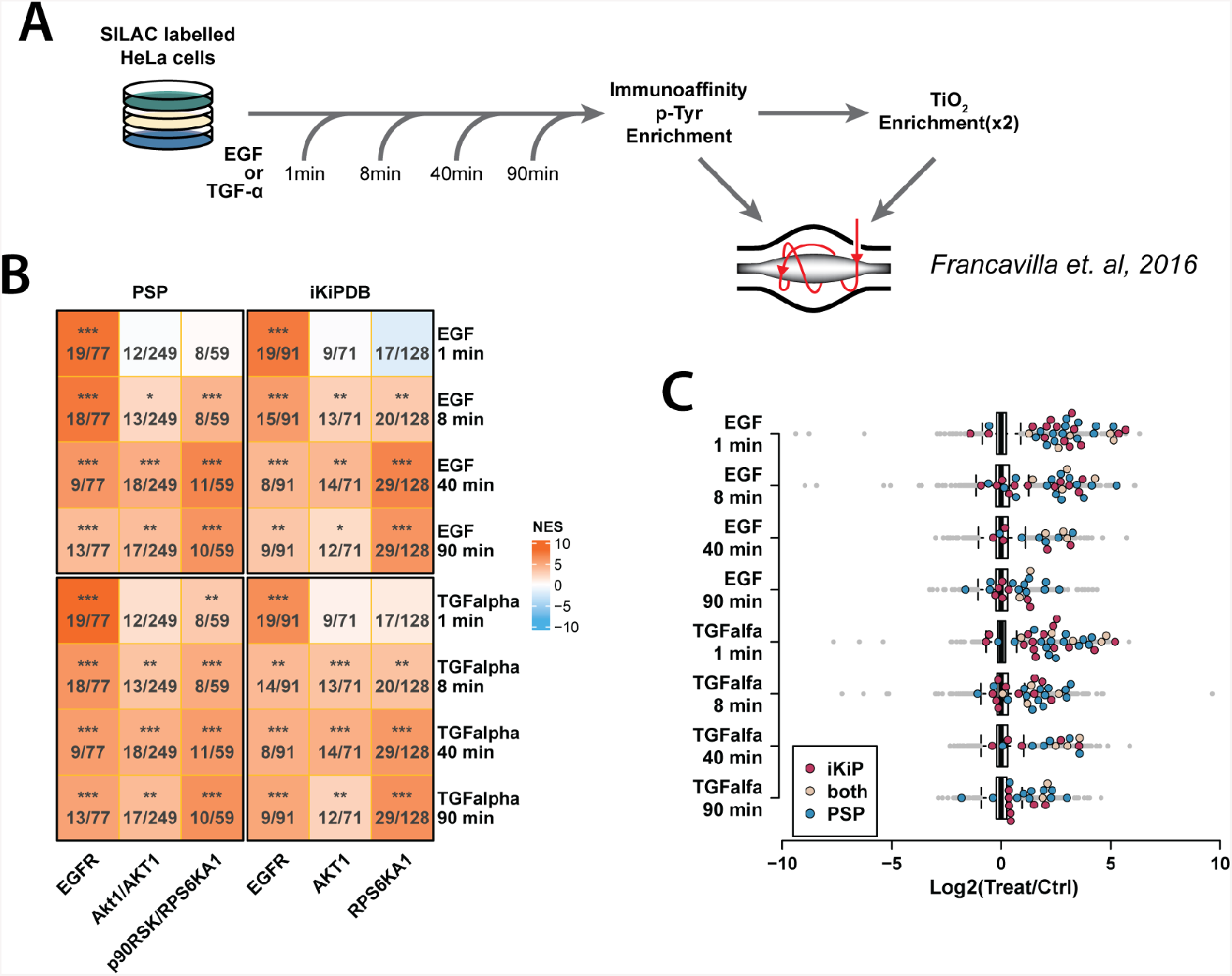
Benchmarking of iKiP-DB. (A) Schematic representation of the experimental design of the data from Francavilla et al. (B) Heatmap depicting the prediction output and colored according to the normalized enrichment score of the PTM-SEA analysis. Each cell reports the significance of the prediction (permutation based p-values: * < 10%, ** < 5%, *** < 1%), as well as the size of the overlap over the size of the specific kinase set. (C) Distribution of phosphosites ratios for EGF and TFGɑ treatments. Sites annotated for EGFR in iKiP-DB, PSP or both are highlighted.

### iKiP-DB predicts the inhibition of well and poorly characterized kinases

To further validate iKiP-DB as a useful tool to predict kinase activity, we searched the literature for phosphoproteomics datasets with the following characteristics: i) treatments with kinase inhibitors or knockdowns, as they would provide an easy ground truth to test our predictions; ii) studies focusing on a range of kinases, both well and less characterized; iii) datasets derived from samples other than HeLa cells, to test predictions across different model systems. According to these criteria, we selected four additional datasets for further benchmarking of iKiP-DB (Figure 3). Again, we used the PTM-SEA algorithm to compute kinase enrichment scores^15^.

**Figure 3.**
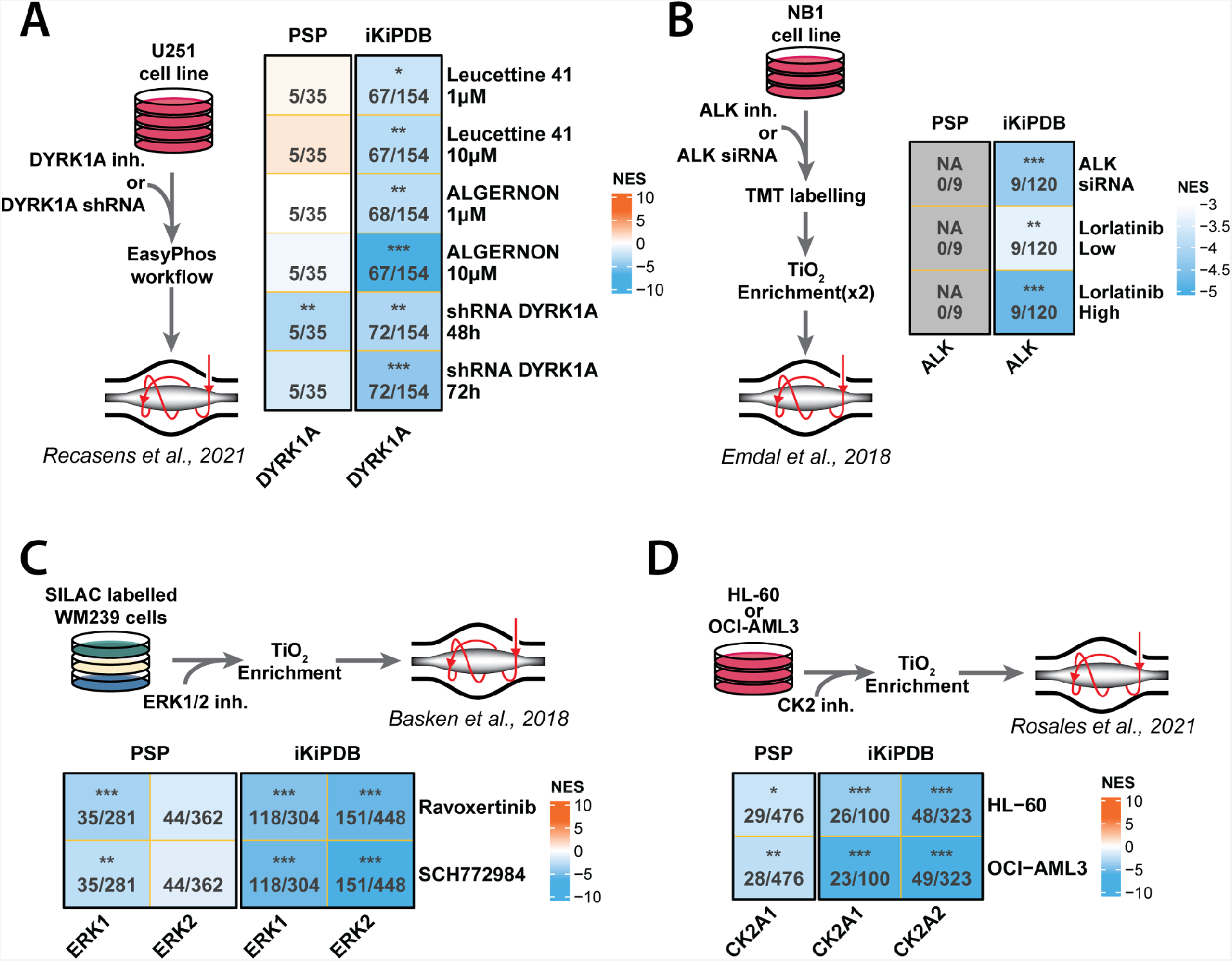
iKiP-DB correctly predicts kinase inhibitions under different experimental conditions. All panels depict the experimental design of the data used on the left or top and a heatmap with the output of the prediction on the right or bottom. Each heatmap is colored according to the normalized enrichment score (NES) and each cells reports the significance (permutation based p-values: * < 10%, ** < 5%, *** < 1%), as well as the size of the overlap over the size of the specific kinase set. We tested our database on the inhibitions of DYRK1A in glioblastoma cells (A), ALK in neuroblastoma cells (B), ERK1/2 in metastatic melanoma cells (C) and CK2 in two acute myeloid leukemia cell lines (D).

For the selected datasets on DYRK1A^22^ and ALK^23^ kinase inhibition, PSP provides poor predictions of activity (Figure 3A, Supplemental Table S3), or no prediction at all because no observed sites overlapped with the database (Figure 3B, Supplemental Table S3). In contrast, the predictions made with iKiP-DB reflected the treatments the cells were subjected to: U251 glioblastoma cells treated with two DYRK1A inhibitors at two different concentrations show an inhibition for the correct kinase, with the normalized enrichment scores (NES) reflecting the increased concentrations of the inhibitors with lower scores. A similar behavior is observed for DYRK1A knockdowns, with lower NES for the longer time point of knockdown, providing an excellent confirmation of the validity of our kinase activity annotation (Figure 3A and S2A). Additionally, iKiP-DB indicated DYRK1A among, when not the most, inhibited kinase (Figure S2B). Similarly, analysis with iKiP confirms the knockdown of ALK in neuroblastoma NB1 cells, as well as a dose-dependent inhibition with lorlatinib (Figure 3B, S2C and S2D).

The next two dataset we selected focused on the inhibition of ERK1/2 and CK2, which are well studied and annotated kinases in PSP (Figure 1E). ERK1/2 inhibitors are of particular interest due to their role in the clinic, for example for the treatment of melanoma, especially after the development of metastases^37^. In their work, Basken and colleagues tested two potent ERK1/2 inhibitors on WM239 metastatic melanoma cells and measured their effect on the phosphoproteome^24^. Kinase activity prediction with iKiP-DB correctly indicated the strong inhibition of both ERK1 and 2, while prediction with PSP could only detect the significant depletion of ERK1 (Figure 3C, Supplemental Table S3). Next, we wanted to test whether our database could be used to unbiasedly identify the correct target of a kinase inhibitor. For this, we assessed which kinases were predicted as most inhibited by both drug treatments. Analysis with iKiP-DB correctly identified ERK2 as the most inhibited kinase by ravoxertinib and SCH772984 (Figure S2E), which is well in line with the lower IC50 values of both compounds for ERK2 compared to ERK1^38,39^. On the contrary, analysis with PSP did not indicate either ERK as the most inhibited kinase by either treatment (Figure S2E, Supplemental Table S3). Finally, in the last study we used for benchmarking, Rosales and colleagues treated two acute myeloid leukemia cell lines with a peptide-based kinase inhibitor targeting CK2^25^. While both PSP and iKiP-DB correctly predicted the inhibition of CK2A1, our database could also provide information regarding the inhibition of CK2A2, the other catalytic subunit of the kinase (Figure 3D, Supplemental Table S3). Again, we assessed which kinases were predicted as most inhibited to evaluate the potential of our database for unbiased target discovery. Also here, iKiP-DB could be used to correctly assess the target of the inhibitor in all conditions (Figure S2F, Supplemental Table S3).

In conclusion, reanalysis of published phosphoproteomic datasets highlights the ability of iKiP-DB to predict the activity of different kinases in various cell lines and with data generated via different quantification and enrichment strategies (SILAC, TMT, EasyPhos, TiO2, anti-p-Tyr antibodies). Hence, iKiP-DB can robustly predict kinase activity in diverse phosphoproteomic datasets.

### iKiP-DB identifies activated kinases upon SARS-CoV-2 infection

As proof of concept, we applied iKiP-DB to a new dataset of SARS-CoV-2 infected lung epithelial cells. Briefly, we infected lung epithelial cells (Calu-3) with SARS-CoV-2 (Patient isolate, BetaCoV/Munich/BavPat1/2020|EPI_ISL_406862) at a multiplicity of infection of 0.33, and collected samples at 4, 8 and 12 hours post infection (hpi) in triplicates, along with mock infected controls. For quantitation and multiplexing we employed isobaric labelling with tandem mass tags (TMTpro)^29,30^. From these samples we acquired total proteome and IMAC-enriched phosphopeptide data (Figure 4A), which yielded over 8000 proteins and approximately 13000 phosphosites with no missing values. While the impact of the infection on host proteome and phosphoproteome was modest (Figure S3), we measured a strong increase of all SARS-CoV-2 proteins, confirming the presence of a productive infection in Calu-3 cells (Figure 4B, Supplemental Table S4). At the latest time point, we identified several differentially regulated host proteins (Figure 4C, Supplemental Table S4). Among these, many interferon-stimulated genes such as IRF7, IFIT1, IFIT2, IFIH1, OAS1 and MX2, were enriched upon infection. These proteins are well characterized for their antiviral function^40,41^, also in the context of SARS-CoV-2 infection^27,42,43^. We also identified downregulation of ACE2, the most important entry factor of SARS-CoV-2^44^ which is in line with the reported shedding of the receptor from infected cells^45^.

**Figure 4.**
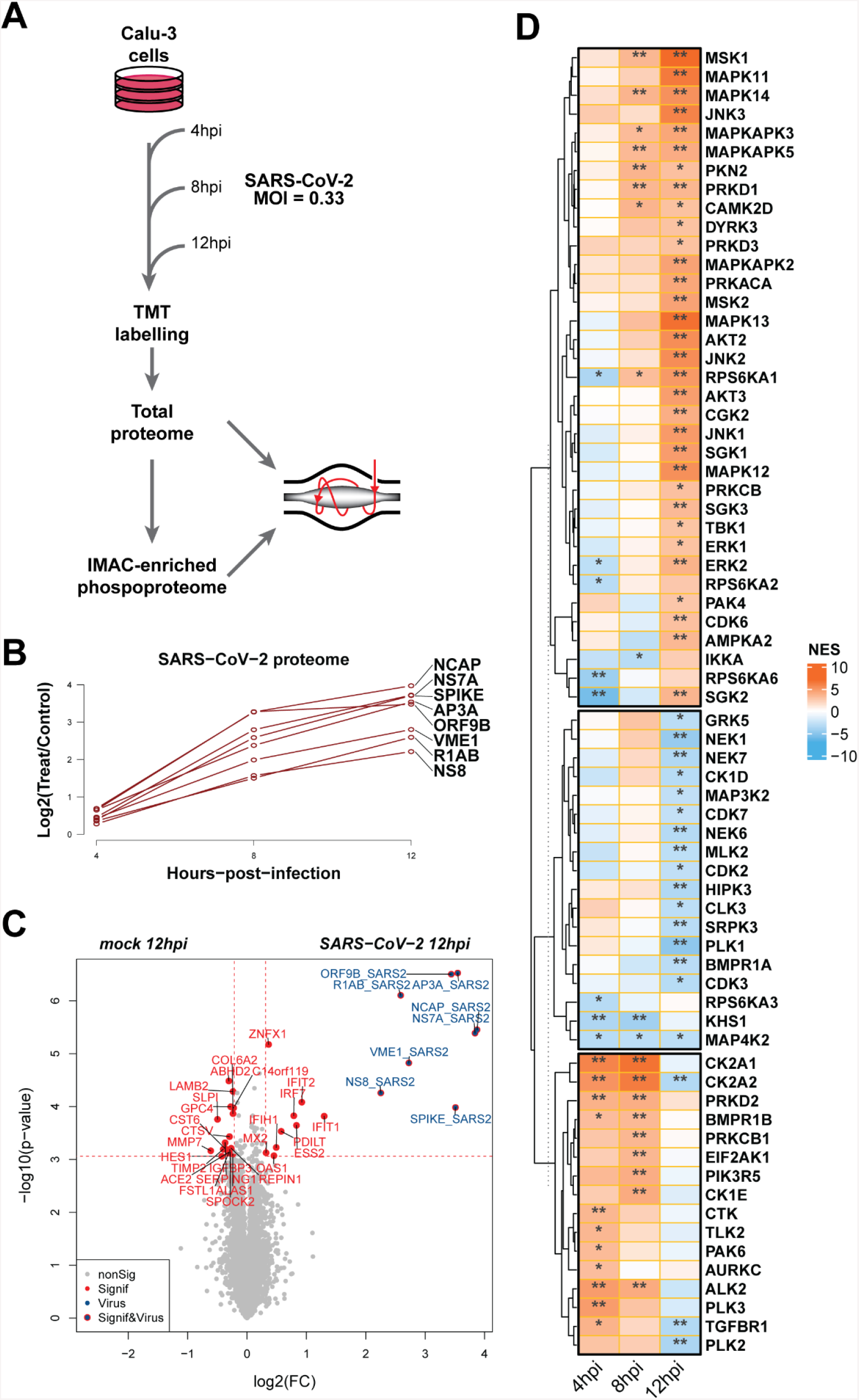
Changes in kinase activities upon infection with SARS-CoV-2. (A) Schematic representation of our infection experiments of lung epithelial cells with SARS-CoV-2. (B) Variation in abundance of all quantified SARS-CoV-2 proteins throughout the course of the infection. (C) Volcano plot depicting the significantly regulated proteins (host and viral) at the latest time point measured. The red lines indicate the cut-off used for significance calling. (D) Variation in kinase activity caused by the infection, as predicted by iKiP-DB. The heatmap is colored based on the normalized enrichment score (NES) of the PTM-SEA analysis, and each cell reports the significance cut-off (Benjamini-Hochberg corrected p-values: * < 10%, ** < 5%, *** < 1%). Kinases were divided into three clusters by k-means clustering based on the NES values.

To gain insight on how the virus changes the activity of host kinases, we analyzed the phosphoproteomic data using iKiP-DB (Figure 4D, Supplemental Table S4). This analysis divided kinases into three broad groups: early activated, late inhibited and late activated, with the last group being the largest (Figure 3C). Of particular interest is the presence of TANK binding kinase 1 (TBK1) among the kinases activated at the latest time point post infection. This is well in accordance with the role of TBK1 as a downstream effector of RIG-I-like receptor signaling^46^ and also with the induction of interferon stimulated genes we observed at 18 hpi. Additionally, many of the late activated kinases are known to be involved in the response to viral infections, such as MAP kinases^47^ and stress-activated kinases downstream of the MAP/ERK pathway like RPS6KA1^48^. Among the kinases predicted to be downregulated, several are involved in cell cycle progression, including Polo-like kinases (PLKs) and NimA related kinases (NEKs)^49,50^, in line with the decreased proliferation induced by the infection. Overall, our analysis is consistent with current knowledge of the effects of SARS-CoV-2 infection at the cellular and phosphoproteome levels^27,51^, reflecting the cellular response to infection.

## CONCLUSIONS

Protein phosphorylation is a fundamental and ubiquitous process in biological systems^52^, and as such studying the phosphoproteome is highly informative for basic biology and disease^21,53^. While novel technologies are rapidly increasing the depth and coverage of MS-derived phosphoproteomics data, their interpretation remains challenging, partially due to the scarcity of functional annotations of phosphosites^6^.

In this work, we described the re-analysis of published *in vitro* kinase assay data, which we compiled into a database of kinase-substrate associations we named iKiP-DB. Compared to PSP as the current gold standard for phosphosite annotation^5^, our database increases the range of annotated kinases as well as the number of substrate sites annotated. Additionally, since our database is derived from experimental data, it is not biased towards the most well-studied portion of the kinome. We employed iKiP-DB in conjunction with PTM-SEA^15^ to predict the activation or inhibition of kinases in published phosphoproteomics data. Direct comparison demonstrated an equal or higher degree of accuracy of iKiP-DB compared to the manually annotated data of PSP. Interestingly, we also observed a superior prediction by iKiP-DB for CK2 and ERK1/2 inhibitor experiments, even though both CK2 and ERK1/2 are well annotated kinases in PSP. Finally, analysis of newly generated data of SARS-CoV-2 infected lung epithelial cells recapitulates the phenotypic effects of viral infections, as well as the known biology of the novel coronavirus.

One important limitation of iKiP-DB is that it is based on *in vitro* kinase experiments, which do not necessarily reflect the activity of kinases *in vivo*. We alleviate this problem by restricting the database to sites that have been observed in cells (Figure 1B). Nevertheless, individual phosphosites in iKiP-DB are not necessarily phosphorylated by the corresponding kinase under physiological conditions. Therefore, iKiP-DB is more useful for the large-scale analysis of kinase signatures, rather than at the level of individual phosphosites. It is actually surprising how well the *in vitro* kinase data alone (without any manual annotation) reflects cellular kinase activity (Figure 2 and 3).

In summary, we demonstrate that integrating phosphoproteomic datasets with *in vitro* kinase data via iKiP-DB greatly facilitates detection of altered kinase activity. We believe this tool will be broadly useful for phosphoproteomic data analysis. We provide iKiP-DB in GMT format, ready to use for PTM-SEA using the ssGSEA suite.

## Supporting information

iKiP-DB in GMT format

## ACKNOWLEDGMENTS

We would like to thank Dr. Evelyn Ramberger for the critical reading of the manuscript, as well as Mohamad Haji of the mass spectrometry core facility of the MDC for his invaluable help in sample preparation. Since this work would have not been possible without all the published data we used for the database and the benchmarking, we would like to express our gratitude to all the researchers involved in these studies that generated high-quality phosphoproteomics data we could use. We would like to especially mention Naoyuki Sugiyama, Yasushi Ishihama and Haruna Imamura, since the core of this work was based on their dataset. This work was supported by the German Ministry of Education and Research (BMBF) via the national research node for mass spectrometry in systems medicine MSTARS (031L0220B to M.S.).

## Notes

### Competing Interest Statement

The authors have declared no competing interest.

