## Supplemental Figures 1-3 for "In vitro Kinase-to-Phosphosite database (iKiP-DB) predicts kinase activity in phosphoproteomic datasets"

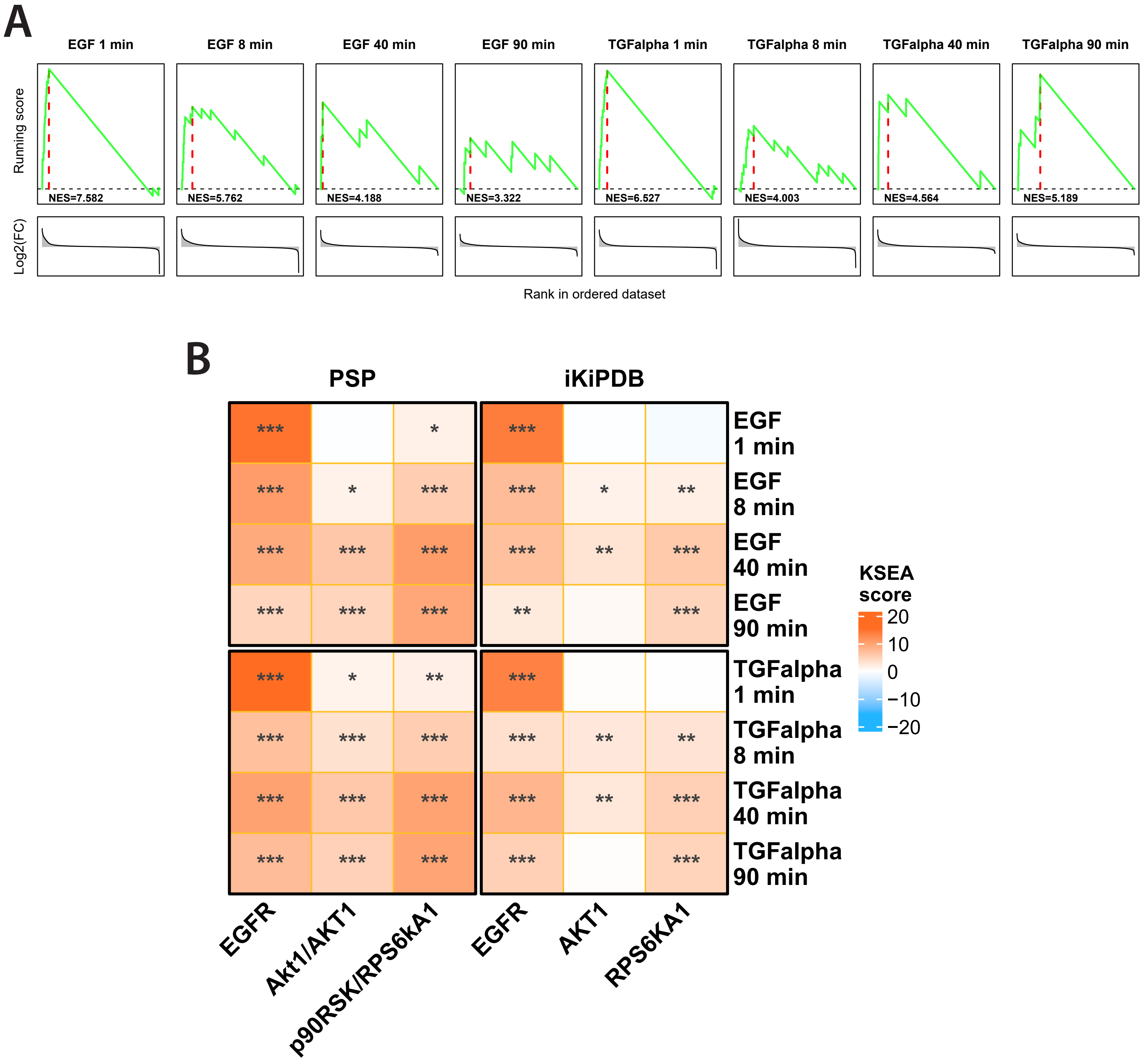


**Supplemental Figure 1**. Relative to Figure 2. (A) Running scores for the PTM-SEA calculation of the EGFR kinase set in iKiP-DB for the experiments shown in Figure 2. The green lines represent the value of the running score at each point of the rank-ordered phosphosites, and the red line is the maximum deviation from 0. NES values are reported for each condition, as well as the full distribution of phosphosites ranked based on log2-transformed treatment over control rations (below). (B) Heatmap depicting the outcome of the KSEA analysis on the same dataset shown in Figure 2. The cells are colored based on the KSEA score and each cell reports the significance of the enrichment (one-tailed p-value: * < 10%, ** < 5%, *** < 1%).

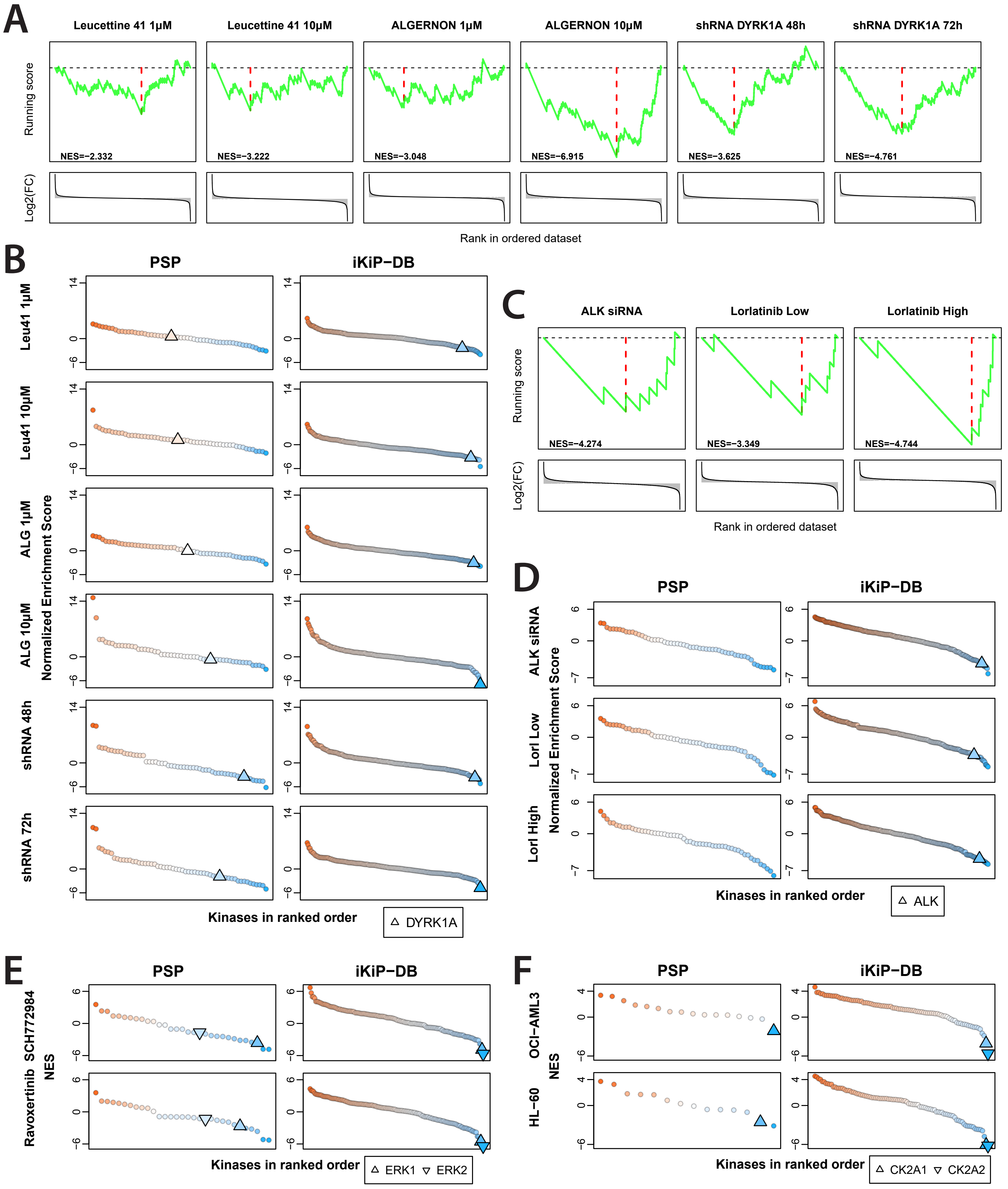


**Supplemental Figure 2**. Relative to Figure 3. (A) and (C) represent the running scores for the PTM-SEA calculation for the experiments shown in Figure 3A and 3B, respectively. The green lines represent the value of the running score at each point of the rank-ordered phosphosites, and the red line is the maximum deviation from 0. NES values are reported for each condition, as well as the full distribution of phosphosites ranked based on log2-transformed treatment over control rations (below). (B), (D) (E) and (F) show the kinases ranked by NES for the experiments analyzed in Figure 3A, 3B, 3C and 3D respectively. All kinase sets with at least 5 phosphosites overlapping with the sites measured are shown, irrespective of the significance of the enrichment. The relevant kinases for each experiment are highlighted.

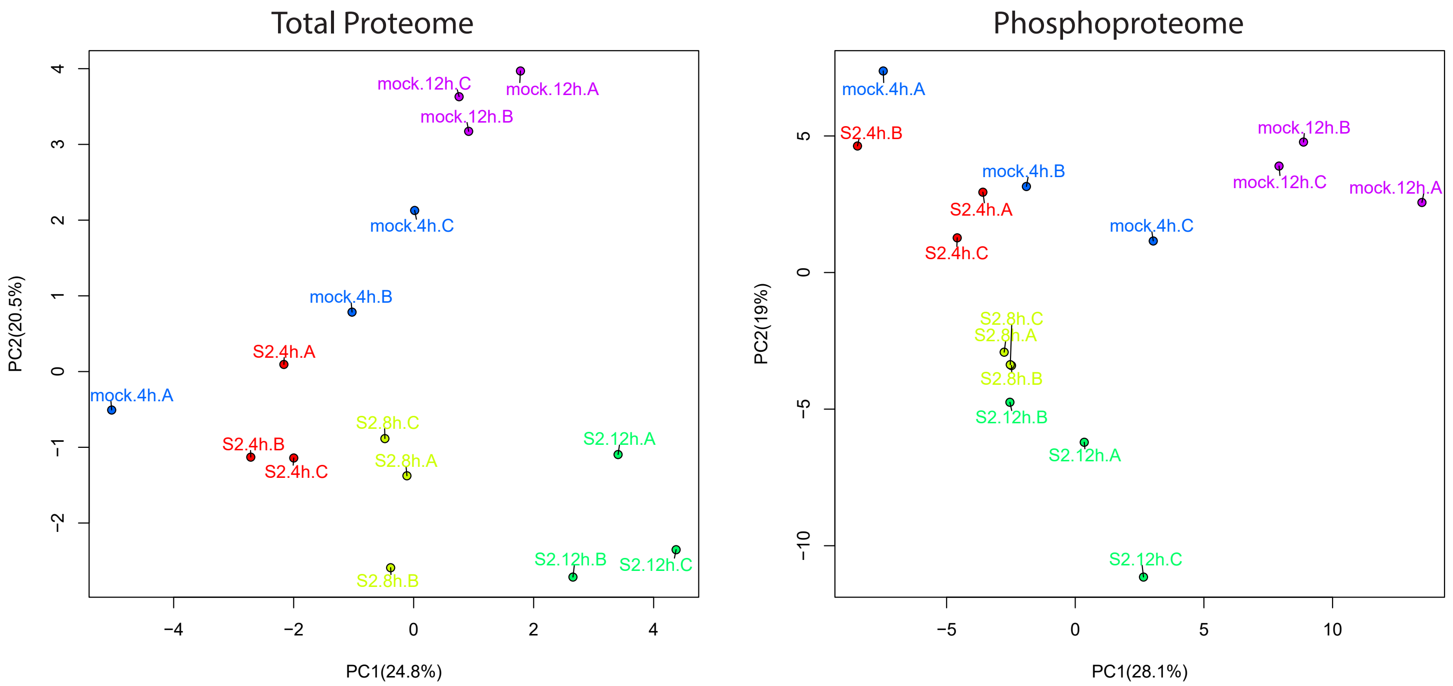


**Supplemental Figure 3**. PCA plots of proteomics and phosphoproteomics measurements of SARS-CoV-2-infected Calu-3 cells, including mock infected controls.
